# Induction of autotetraploidy and microbiome associations mediate differential responses to pathogens

**DOI:** 10.1101/2021.10.01.462589

**Authors:** Elijah C Mehlferber, Michael J Song, Julianne Naomi Pelaez, Johan Jaenisch, Jeremy E Coate, Britt Koskella, Carl J Rothfels

## Abstract

It has become increasingly clear that the microbiome plays a critical role in shaping the host organism’s response to disease. There also exists mounting evidence that an organism’s ploidy level is important in their response to pathogens and parasites. However, no study has determined if or how these two factors influence one another. We investigate the effect of whole-genome duplication in *Arabidopsis thaliana* on their above-ground (phyllosphere) microbiome, and determine the interacting impacts of ploidy and the microbiome on disease outcome. Using seven independently derived synthetic auto-tetraploid *Arabidopsis* accessions, a synthetic leaf-associated bacterial community, and the model pathogen *Pseudomonas syringae* pv. Tomato DC3000, we confirm that polyploids are generally more resistant to pathogens, but illustrate that this resistance may be in part due to a decrease in the reliance on beneficial bacteria. Polyploids fare better against the pathogen than diploids regardless of microbial inoculation, while we observed that diploids harboring an intact microbiome have lower pathogen densities than those without. We then use RNA sequencing to show that diploids have many more differentially expressed defense-related genes in the presence of their phyllosphere microbiota, while polyploids exhibit constitutively activated defenses regardless of exposure to the synthetic community. These results imply that whole-genome duplication can disrupt historical host-microbiome associations, and suggest that a potential cause or consequence of disruption is a heightened capacity for pathogen defense that is less impacted by the microbiome.

## Introduction

Whole-genome duplications (WGDs), or “polyploidizations”, are dramatic evolutionary events where the entire genome is doubled, followed by a period of gene silencing and neo-funcionalizion that can lead to the extension or divergence of ecological niches from the parent range (Hijmans et al., 2007; Theodoridis et al., 2013; Molina-Henao and Hopkins, 2019) and are often considered to be drivers of evolution (reviewed in Van de Peer et al., 2017). Polyploidy is associated with many novel and potentially adaptive phenotypes including changes to biomass, photosynthesis, water- and nitrogen-use efficiency, and secondary metabolism (Ni et al., 2009; Coate et al., 2012; Huang et al., 2007; Levin, 1983), with polyploids having larger cells and organs and more chloroplasts per cell (Coate et al., 2012). To this end, polyploidy is often considered to be a potential mechanism by which short-term adaptations may arise in response to changes to the environment or stress (reviewed in Van de Peer et al., 2017). Furthermore, theoretical models predict that increases in ploidy will increase resistance to parasites and pathogens (Oswald and Nuismer, 2007), and there is some experimental evidence that supports this conclusion. For example, in *Actinidia chinensis* (kiwifruit), hexaploids are the most resistant to pathogenic *Pseudomonas syringae*, followed by tetraploids and then diploids (Saei et al., 2017), and inducing polyploidy in *Impatiens walleriana* (cultivated impatiens) confers increased resistance to mildew (Wang et al., 2018). This phenomenon of increased resistance parallels the ability of the plant’s associated microbial community (the microbiome) to also play a critical role in the plant’s defense against pathogens (Wei et al., 2019; Leopold and Busby 2020).

Both the root and shoot systems of plants host diverse microbial communities, including bacteria, fungi, and other eukaryotes, but these plant systems associate with only a subset of all environmentally available microbes, many of whom play important functions in disease or nutrient acquisition (reviewed in Bulgarelli et al., 2013). Which taxa successfully colonize a given plant can be mediated by the host both directly and indirectly, including through direct immune responses (Lebeis et al., 2015), coordination of stress and immune system functions (Castrillo et al., 2017), or the production of secondary chemicals (Levin, 1983). Historically, the study of plant-microbe interactions has focused on the plant response to pathogens (Tao et al., 2003) and on identifying the genes involved in defense responses (Mahalingam et al., 2003; Zhu et al., 2013). Recently, this interest has broadened to include the role that commensal and mutualistic bacteria, which comprise the majority of plant-associated microbes, play in the functioning of their host. Host reliance on the microbiome for disease resistance has recently been considered as a key determinant of immune system evolution (King & Bonsall 2017; Metcalf & Koskella 2019; McLaren et al. 2020), and thus ploidy-induced changes in microbiome-mediated defense could have important consequences for subsequent host evolution. As such, a current open, yet critical, question is how a whole-genome duplication event might impact the interaction between plants and these associated microbiota.

In order to determine how WGDs might alter the interactions between the plant and its above-ground microbiota and/or pathogen growth, we used seven lines of synthetic auto-tetraploid accessions of *Arabidopsis thaliana*, and, along with their diploid progenitors, inoculated them with a synthetic community (SynCom) comprising taxa common to the leaf habitat. We then determined whether there was a conserved change in bacterial community composition across ploidy level, and if plants of differing ploidy had different transcriptional responses to these bacteria. To investigate any effects of these changes on microbiome-mediated pathogen protection, we inoculated these plants, along with untreated controls (plants without the synthetic microbiome), with the model *Arabidopsis* pathogen *Pseudomonas syringae* pv. tomato DC3000 and measured growth during early establishment. By using synthetic auto-tetraploid accessions of *Arabidopsis* in conjunction with a controlled, synthetic microbial community, we were able to assess the associations between genotype, ploidy level, and the microbiome, determine the extent to which these interactions are mediated through shared transcriptional responses, and quantify the effect of these interactions on pathogen defense.

## Results

### Ploidy level does not impact microbiome establishment

Alpha diversity of the established microbiome (measured as species richness, Shannon index, or species evenness) did not differ significantly between diploids and polyploids (pairwise ANOVA, P>0.05; Figure 1 A & B). Likewise, we found no significant differences in beta diversity (community composition measured with Bray-Curtis dissimilarity) across ploidy level (ADONIS nonparametric multivariate analysis of variance, P>0.05; Figure 1 C).

**Fig. 1.**
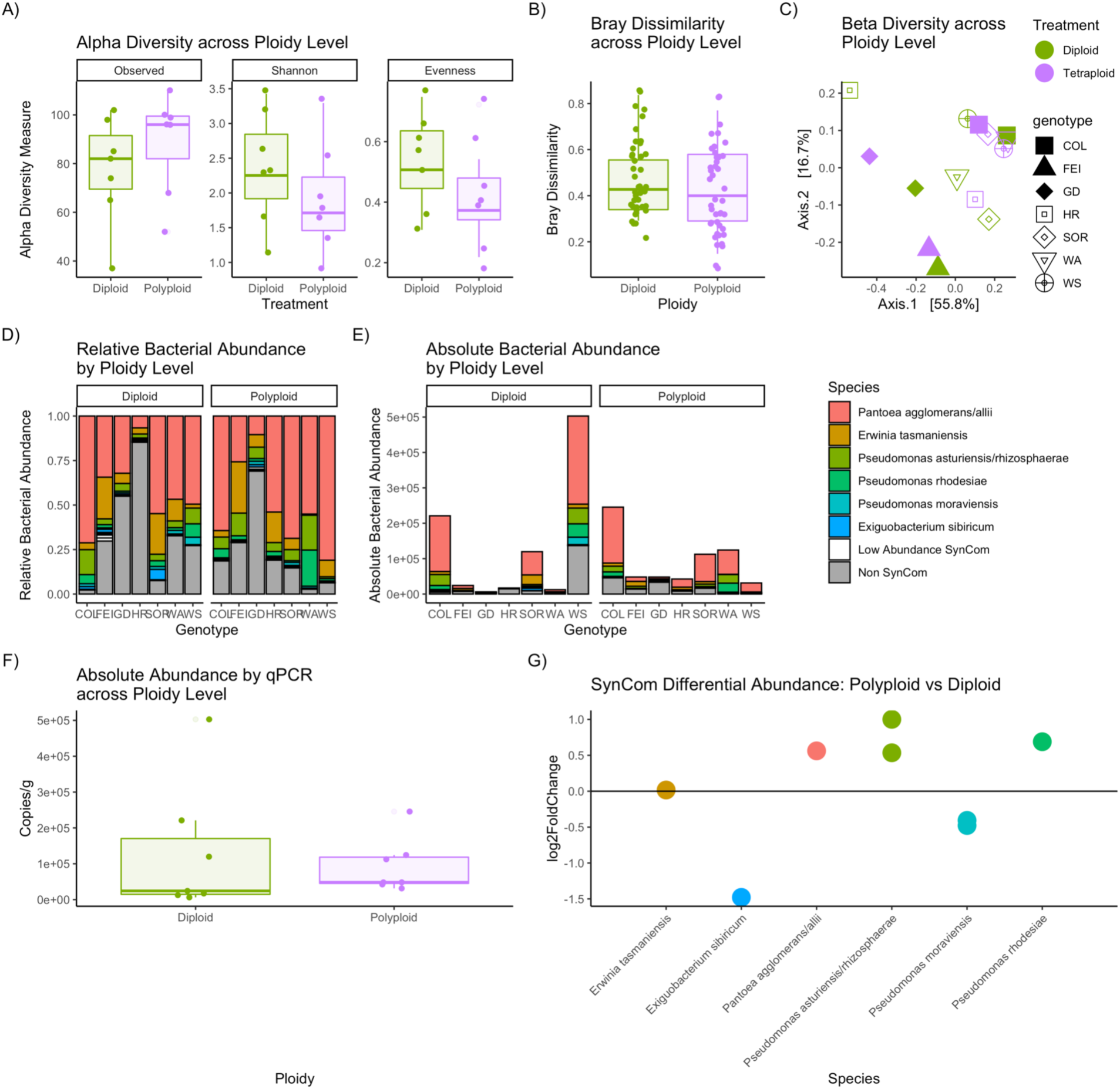
The effect of ploidy on microbiome composition and structure. A) There is no significant difference in the tested Alpha Diversity metrics, inducing Observed, Shannon, and Eveness. However, there is a non-significant trend towards lower Shannon diversity in the Polyploid plants, which is driven primarily by their lower Eveness. B) Bray-Curtis dissimilarity between plants of each treatment is not significantly different. C) We visualize Bray-Curtis dissimilarity across treatments using a PCoA plot finding that there is no significant grouping by treatment. D) A bar graph illustrating the relative abundance of the SynCom inoculated ASVs across treatments, as well as the residual community. E) A bar graph visualizing the qPCR derived absolute abundance of the microbial communities across treatments. F) There is no significant difference in the qPCR derived absolute abundance of phyllosphere bacteria between the two treatments. G) A plot showing the differential abundance of SynCom ASVs between the polyploids and diploids, despite some differences in abundance these differences are not significant.

As expected, the vast majority of bacteria that we found associated with the plants were from the synthetic community, with *Pantoea, Pseudomonas*, and *Exiguobacterium* showing consistently high relative abundance across samples (Figure 1 D,E). qPCR analysis suggests that the absolute abundance of bacteria from the SynCom on the leaves one week after inoculation was not significantly different across ploidy levels (Figure 1 F; standardizing for sample weight (Supplemental Figure 2): Welch Two-Sample t-test, t= -0.11455, df = 10.076, p = 0.911). There were increased numbers of *Bacillus* and *Frigobacterium* on tetraploids versus diploids, as well as decreased abundance of *Exiguobacterium* and *Lysinbacillus* in the tetraploids, but these patterns were not significant (Figure 1 G, Supplemental File 1).

### Tetraploids are less susceptible to pathogen establishment

Time since exposure to the pathogen and the presence of a microbiome both significantly impacted the pathogen abundance (Linear Mixed Effects Model, p = 0.0018 and p = 0.0031, respectively; Table 2), and we found a marginally significant interaction between the two (p = 0.0517; Table 2). For the diploid samples analyzed alone we found a significant impact of time, treatment, and their interaction (p = 0.0001, p = 0.0003, p = 0.01, respectively; Table 2). We performed a post-hoc test (Tukey HSD), finding significant differences between SynCom-treated samples across timepoints one and two (p = 0.0008), between SynCom-treated samples from timepoint one and control samples from time point 2 (p = 0.0003), and between SynCom-treated and control samples in time point 1 (p = 0.0014). We determined that it was inappropriate to evaluate the polyploid samples, as the addition of any terms to the model showed no significant improvement over a null model including only the intercept. In addition, there was significantly lower pathogen abundance on the polyploid plants at the second time point for control plants (Welch Two-Sample t-test, t = 2.809, df = 4.9939, p = 0.03765), as well as treated plants (Welch Two-Sample t-test, t = 2.4295, df = 8.211, p = 0.04048; Figure 3C).

**Table 1:** Linear Mixed Effects Model (nlme) of DC3000 Abundance as a function of the explanatory variables Time, Ploidy, Treatment, and their interactions.

| **DC3000 Abundance (Log10)** |  |  |  |
| --- | --- | --- | --- |
| *Predictors* | *Estimates* | *CI* | *p* |
| (Intercept) | 1.15 | 0.18 – 2.11 | **0.033** |
| time [2] | 2.34 | 1.03 – 3.64 | **0.002** |
| ploidy [4] | 0.06 | -1.25 – 1.36 | 0.935 |
| treatment [C] | 2.20 | 0.89 – 3.51 | **0.003** |
| time [2] * ploidy [4] | -1.73 | -3.59 – 0.12 | 0.088 |
| time [2] * treatment [C] | -1.99 | -3.83 – -0.14 | 0.052 |
| ploidy [4] * treatment [C] | -1.40 | -3.25 – 0.44 | 0.164 |
| (time [2] * ploidy [4]) * treatment [C] | 0.55 | -2.13 – 3.24 | 0.702 |
| **Random Effects** |  |  |  |
| $\sigma^2$ | 1.46 | | |
| $\tau_{00 \text{ eco}}$ | 0.14 | | |
| ICC | 0.09 |  |  |
| $N_{\text{eco}}$ | 7 | | |
| Observations | 54 |  |  |
| Marginal R<sup>2</sup> / Conditional R<sup>2</sup> | 0.404 / 0.456 |  |  |

**Table 2:**
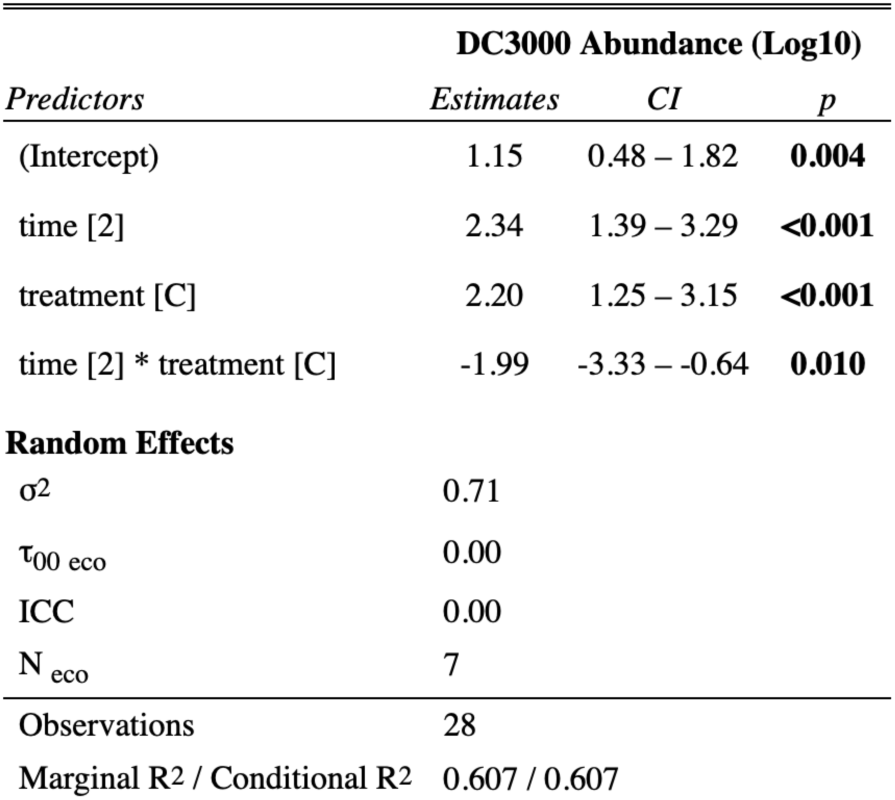
Linear Mixed Effects Model (nlme) for Diploids, with DC3000 Abundance as a function of the explanatory variables Time, Treatment, and their Interactions

| **DC3000 Abundance (Log10)** |  |  |  |
| --- | --- | --- | --- |
| *Predictors* | *Estimates* | *CI* | *p* |
| (Intercept) | 1.15 | 0.48 – 1.82 | **0.004** |
| time [2] | 2.34 | 1.39 – 3.29 | **<0.001** |
| treatment [C] | 2.20 | 1.25 – 3.15 | **<0.001** |
| time [2] * treatment [C] | -1.99 | -3.33 – -0.64 | **0.010** |
| **Random Effects** |  |  |  |
| $\sigma^2$ | 0.71 | | |
| $\tau_{00 \text{ eco}}$ | 0.00 | | |
| ICC | 0.00 |  |  |
| $N_{\text{eco}}$ | 7 | | |
| Observations | 28 |  |  |
| Marginal R<sup>2</sup> / Conditional R<sup>2</sup> | 0.607 / 0.607 |  |  |

**Table 3:** Post hoc Tukey HSD (emmeans) for the Diploid Linear Mixed Effects Model

| *contrast* | *estimate* | *SE* | *df* | *t.ratio* | *p.value* |
| --- | --- | --- | --- | --- | --- |
| 1 B - 2 B | -2.34 | 0.49 | 18 | -4.79 | 0.00 |
| 1 B - 1 C | -2.20 | 0.49 | 18 | -4.51 | 0.00 |
| 1 B - 2 C | -2.55 | 0.49 | 18 | -5.24 | 0.00 |
| 2 B - 1 C | 0.14 | 0.49 | 18 | 0.28 | 0.99 |
| 2 B - 2 C | -0.21 | 0.49 | 18 | -0.44 | 0.97 |
| 1 C - 2 C | -0.35 | 0.49 | 18 | -0.72 | 0.89 |

### Diploid plants exhibit greater response to synthetic community colonization while Polyploids activate defense responses regardless

Analyzing RNA sequences from diploid plants prior to pathogen inoculation, using DESeq2, we found 99 up- or down-regulated genes between SynCom treated and untreated plants, while tetraploid plants, again, prior to pathogen exposure, only had 17 significantly differentially expressed genes at the 0.05 p-value cut-off (Fig2; A, B). In particular, diploid plants showed several clusters of significantly differentially expressed genes when those genes were grouped by function. Many of these groups of genes are associated with defense functions. For example, genes associated with the well-characterized phytohormone abscisic acid (ABA) were both up- and down-regulated (Figure 3; A). Genes associated with hypoxia as well as defense response to bacteria were also significantly up- and down-regulated. Furthermore, several genes associated with ethylene signaling, which is integral to the regulation of the plant immune response (Ecker and Davis, 1987), were up-regulated in the SynCom treated diploids but not in the corresponding tetraploids (Figure 3; A). For both microbiome-treated and untreated plants, when compared to the diploids, the polyploids showed a pattern of elevated gene expression across most genes that were differentially expressed (Figure 2; C, D). These genes with elevated expression included defense-related genes, with treated tetraploids having higher expression of ABA, hypoxia, and ethylene signaling related genes relative to untreated tetraploids. In addition, when compared to the diploids, polyploids generally show a pattern of increased defense-gene expression, regardless of their microbiome treatment. Furthermore, we saw fewer significant differences in gene expression in the treated vs. control polyploids, indicating that many of these genes are constitutively expressed. In the cases where we did see significant differences in gene expression, the genes primarily showed increased expression. This is in contrast to the diploids, where genes were modulated both up and down in expression as a result of microbiome treatment.

**Fig 2.**
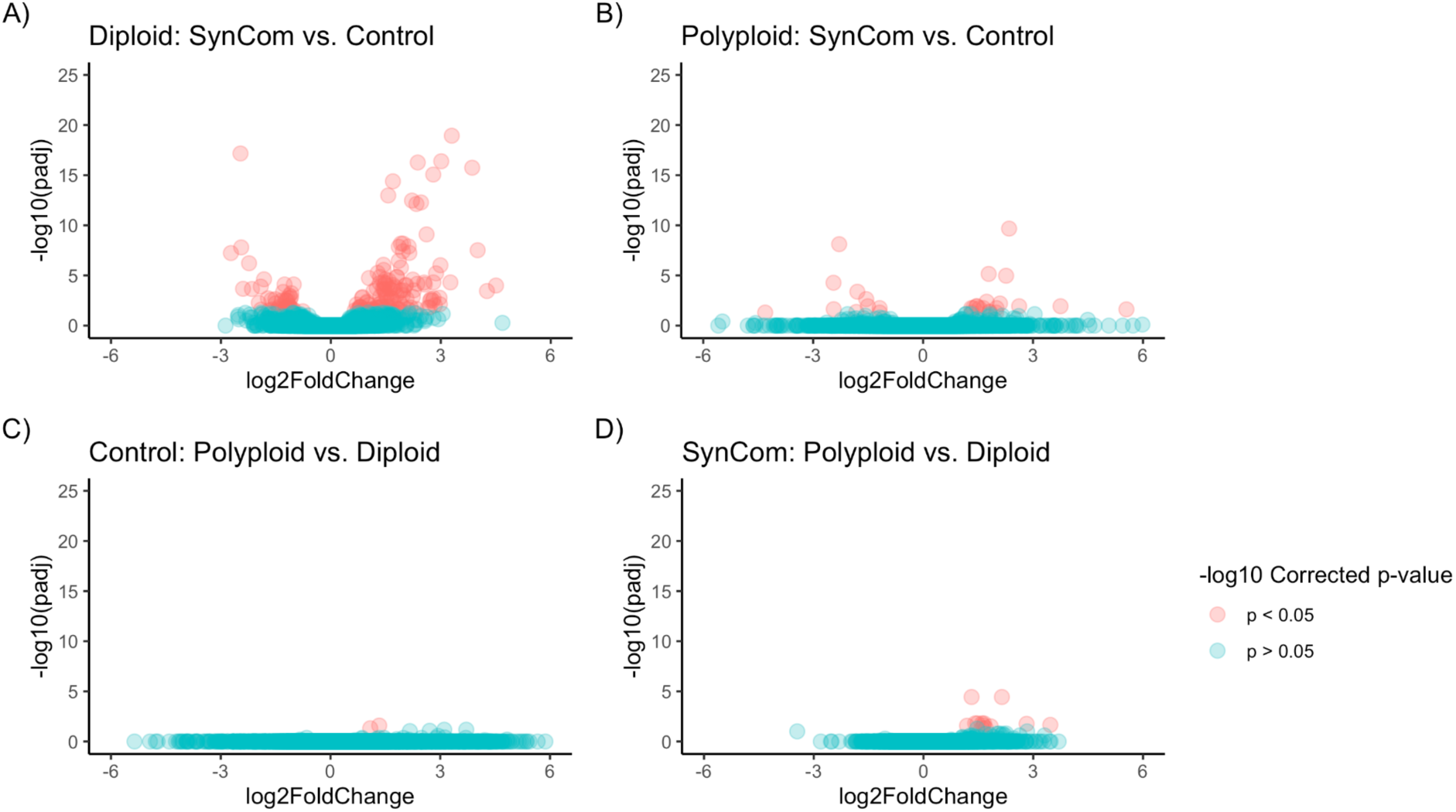
Proportion of total genes measured that reached statistical significance. A-D) Volcano plots showing proportion of total genes that are differentially expressed at the level of p < 0.05 plotted against their log2Fold changes. Comparisons are drawn across Ploidy (Diploid vs. Polyploid) and SynCom treatment (Control vs. SynCom).

**Fig 3.**
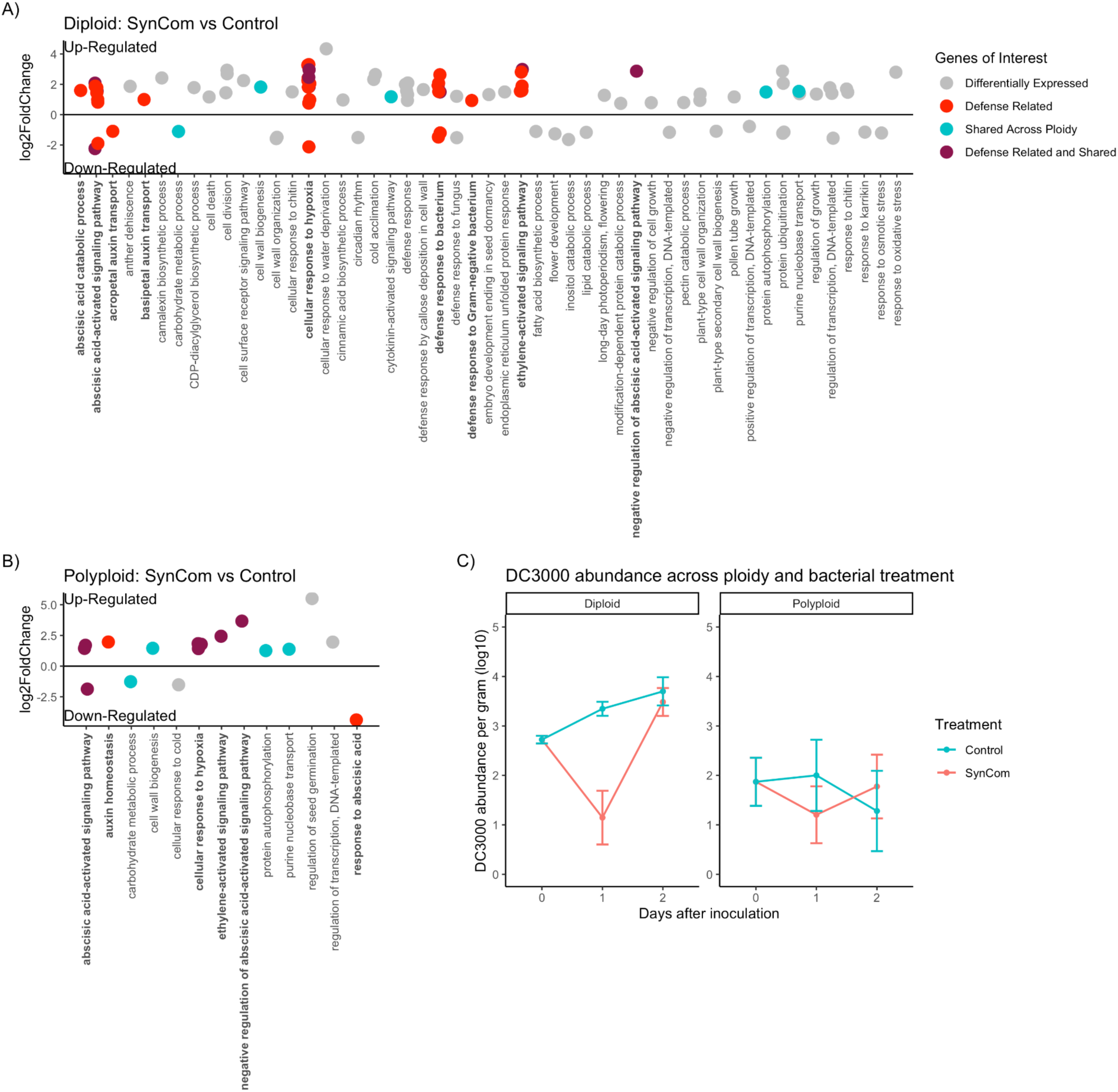
Differential expression of genes highlighted by association with defense response. A) We used Deseq2 to identify genes that are differentially expressed between the microbiome and Control treated Diploids at the level of p < 0.05. Genes that are associated with defense functions are noted in bold and highlighted in red, while genes that are also differentially expressed in the Polyploids are highlighted in blue, and genes that are both defense associated and shared are highlighted in purple. B) Deseq2 informed differential expression between microbiome and Control Polyploids, using the same colors from the previous panel. C) We quantified the abundance of the pathogen DC3000 across timepoints in and treatments in Diploid and Polyploid plants using ddPCR.

## Discussion

### Effects of Polyploidy on Microbiome Diversity

We tested for a conserved shift in the composition of the microbiome as a result of polyploidization by comparing alpha and beta diversity of the phyllosphere across synthetic auto-tetraploid *Arabidopsis* accessions inoculated with a synthetic microbiome. Overall, we found no significant differences in composition or diversity across ploidy levels. The possibility remains that there exist ploidy-dependent differences within a given genotype, which would have been obscured by our experimental design, or that this result is due to the use of a simplified synthetic community. However, our use of multiple accessions allowed us to rule out a generalized response of the microbiome in this system.

Polyploid plants tend to have greater biomass (Pacey et al., 2020), and indeed the polyploid plants in our study weighed significantly more across all accessions. This is in contrast to previous work on autotetraploid *Arabidopsis* (Chen, 2010; Ng et al., 2012). This increased biomass meant that polyploids supported a higher total density of commensal bacteria, but once we normalized for the weight of the plant, these differences were not significant. Likewise, we did not find any significant differences in the relative abundance of any of the synthetic community members across the two ploidy levels, both of which were dominated by *Pantoea, Pseudomonas*, and *Exiguobacterium*. This is in line with work on wheat, where ploidy was found to play a weak and inconsistent role in shaping the below-ground microbiome (Wipf & Coleman-Derr 2021), but contrasts with previous work on *Arabidopsis* which did find a signature of ploidy in shaping microbial community composition across accessions (Ponsford et al. 2020).

### Effects of Ploidy on Pathogen Response

To date there has been very little empirical evidence for a general effect of polyploidy on pathogen response. Although polyploids have been theorized to be more resistant to pathogens (Levin, 1983; Oswald and Nuismer, 2007), empirical studies have generally been inconclusive, i.e. finding evidence for both increased resistance and increased susceptibility (Schoen et al., 1992; Nuismer and Thompson, 2001). Our study leveraged different accessions of *Arabidopsis* in order to discern general patterns between ploidy level and pathogen defense. Overall, we found a trend towards lower pathogen abundance in the polyploid plants regardless of association with a bacterial community, as well as a significant decrease in the abundance of the pathogen in the second time point (Figure 2C).

Autopolyploids (when the genome of one species doubles, such as the *Arabidopsis* used in this experiment) may be more resistant than diploids due to an upregulation of defense genes (King et al., 2012). For example, tetraploid *Arabidopsis* accessions acquired increased resistance to copper stress by having increased activation of antioxidative defense (Li et al., 2017). A buttressing of the antioxidant defense system was also found in colchicine-induced tetraploid plants of *Dioscorea zingiberensis* where antioxidant enzymes were over-produced and maintained at high concentration (Zhang et al., 2010). This generally comes with a fitness trade-off, which was explained in Ng et al. (2012) where they found that proteins associated with stimuli or stress responses were enriched in *A. thaliana* autotetraploids, and that the expression of these genes is associated with a fitness cost and slowed growth. In contrast, our autotetraploids both exhibit greater biomass (Supplemental Figure 3) and defensive capacity (Figure 2), which contradicts this general pattern. Though autopolyploids have increased copy numbers for all genes, polyploidy may have a more significant impact on certain pathways that are involved in both growth and defense which could circumvent the trade offs that are driven by antagonistic crosstalk. This is not to say that our plants do not experience any trade off but at least that this trade off may exist in some metric that we did not collect for the present study, or that they were better at coping with the conditions they experienced, and would manifest the trade off in a less favorable environment.

### Interaction between Ploidy and the Microbiome on Pathogen Response

When assessing the effectiveness of the microbiome in protecting the plants of different ploidy levels, we found that the microbiome temporarily arrests pathogen growth on the diploids, while polyploids are protected regardless of exposure to their microbiome (Figure 2C). This result is particularly interesting in light of previous work on microbiome-mediated protection by a synthetic microbiome in Tomato in which the phyllosphere microbiome was protective against pathogen growth in the absence of fertilizer application to soil, but unimportant when plants had been fertilized prior to microbiome and/or pathogen inoculation (Berg and Koskella, 2018). The plant response to commensal bacterial organisms is complicated, often showing an overlap with the response to pathogens (Vogel et al., 2016). This can be explained in part through the broadly conserved plant responses to common microbial-associated molecular patterns (MAMPs), such as flagellin (Felix et al., 1999), though even these responses can be modulated by a host of commensal interactions, such as repression of conserved epitopes (Colaianni et al., 2021). These responses can be beneficial, through the early activation of broad defense responses (priming) that will then respond more effectively to pathogen exposure (Selosse et al., 2014; Wang et al., 2021). It is possible that this phenomenon plays a role in the increased protection afforded to the diploid plants that have been inoculated with the SynCom, as it may provide a mechanism for them to more effectively prepare themselves for potential future pathogens.

### Genotype drives main transcriptional differences

One of the major challenges in discerning interactions between host and microbiome is the complexity of naturally associated microbial communities. In order to reliably determine the effects of ploidy on microbiome composition, we used a simplified synthetic community, which allowed us to ensure that each plant was exposed to the same bacteria at the same densities at the beginning of the experiment. As expected, the vast majority of the sequences associated with our treated samples were from the inoculum, and the inoculation was broadly successful at establishing communities of known complexity and in high abundance on the plants. This allowed us to assess the transcriptional responses of plants that either had or had not been inoculated with the synthetic community prior to exposure to the pathogen in order to look for general and/or microbiome or ploidy-dependent responses to their commensal microbiome. We found that the broad-scale transcriptional profiles of the samples grouped strongly together by accession (Supplemental Figure 2). This was perhaps unsurprising given that *Arabidopsis* accessions have evolved to be locally adapted to many different environments and have considerable genotypic and phenotypic variation.

### Both polyploids and diploids modulate defense pathways in response to inoculation with the synthetic community

Overall, we found a consistent pattern in the types of gene functions that were differentially expressed in those plants treated with the synthetic microbiome compared to the controls (Figure 2 A-B); most notably, with a host of defense-associated genes showing changes in expression. These genes include those associated with ABA regulation, response to hypoxia, general defense response, and ethylene signaling. ABA is a well-studied plant signaling hormone that is linked to a variety of processes ranging from plant growth, to development and stress response (reviewed in Yoshida et al., 2019). The function of ABA in defense response is multifaceted, and has been shown to be important in pre-invasion defense, through the closing of stomata in response to MAMPs (Melotto et al., 2006), as well as negatively regulating post-invasion defense through the suppression of callose deposition (Clay et al., 2009) and SA dependent resistance (Yasuda et al., 2008), (reviewed in Ton et al. 2009). For all plants that received the synthetic microbiome expression responses were also significantly enriched for GO terms associated with cellular response to hypoxia when compared to the control group. The response to hypoxia requires the ethylene pathway in plants (Fukao and Bailey-Serres, 2004) which is involved in the hormonal control of programmed cell death (Overmyer et al., 2003) and has been shown to influence the composition of the leaf microbial community (Bodenhausen et al., 2014). It also has been found that the response to pathogens involves increased respiration which creates local hypoxia around the leaf which is otherwise fully aerobic (Valeri et al., 2020). Similarly, alcohol dehydrogenase, which in plants is involved in fermentation to produce NAD+, is not only over-expressed in times of low oxygen, but is also induced in response to biotic and abiotic stress and improves responses to pathogens (Shi et al., 2017). Finally, all synthetic microbiome-treated plants showed differential expression for several WRKY transcription factors that are linked with defense signaling (Eulgem and Somssich, 2007), as well as CCR4-associated factor 1, which has been shown to play a role in susceptibility to *P. syringae* infection (Liang et al., 2009). SynCom-treated plants also show a pattern of increased expression in ethylene-activated signaling pathways. Ethylene is another well characterized phytohormone that is responsible for regulation of plant growth, development and senescence (reviewed in Iqbal et al. 2017), as well as response to pathogen invasion and modulation of defense response (Ecker and Davis 1987).

### Polyploids Constitutively Maintain more Defense than Diploids

The trade-offs between growth and defense in plants are often metabolic, but can also be due to the negative interactions between hormones involved in both processes (Karasov et al., 2017). One way to mitigate this tradeoff could be through outsourcing of pathogen defenses to plant-associated microbiota (Karasov et al., 2017). We found that polyploids, regardless of treatment with a commensal microbiome, maintained more defense expression than diploids, with most defense related genes being up regulated. In total, we saw a greater range of differential gene expression in the polyploids as compared to the diploids (Figure 2; B, C, D), indicating that the independent genome duplication events may have had different effects on gene expression among different lines. However, when gene expression reached significance, the majority of those genes were defense related, indicating a conserved response in these genes after the WGD. In contrast, the diploids showed nearly six times the number of differentially expressed genes in total, and a pattern of up and down regulation of defense associated genes. This makes sense in light of our pathogen growth results, as the polyploids are broadly protected regardless of their microbiome, while the diploids seem to require the microbiome to resist pathogen growth. With respect to bacterial pathogen response, diploids significantly downregulated ATCAF1B (Supplemental Figure 4) in the presence of the synthetic community, but polyploids maintained a similar expression profile to the untreated samples. ATCAF1B has been found to be involved with multi-stress resistance (biotic and abiotic) (Walley et al., 2010), in particular in resistance to the pathogen *Pseudomonas syringae* (Liang et al., 2009). One possible explanation for this pattern could be that polyploids are less responsive to the microbiome due to a disruption of fine-tuned pathways as a consequence of a doubling of gene dosage across the genome. When we looked at the *A. thaliana* general non-self response (GNSR) genes (Maier et al. 2021), which see conserved expression changes in the presence of different bacteria, we found that 7 out of the 24 were constitutively expressed at higher levels in the polyploids, regardless of their exposure to the commensal microbiome. Likewise, when comparing the untreated polyploids with the untreated diploids more broadly, we found similar patterns of increased polyploid defense transcription relative to diploids as when we compared treated polyploids with treated diploids.

## Conclusion

Our work highlights the important role that polyploidization plays in mediating the interplay between plants, their associated phyllosphere microbiota and an invading foliar pathogen. While the presence of the synthetic phyllosphere microbiome was always associated with a pattern of decreased *P. syringae* growth, the effect was only significant in the diploid plants, while the tetraploids appeared to be broadly protected regardless of the presence of these beneficial bacteria. Our transcriptional results suggest this is due to a more constitutive, microbiome-independent upregulation of defense genes in the polyploids. It is possible that, as a consequence of gene dosage doubling due to WGD, polyploids are generally more capable of mounting effective defense responses, and may have a higher baseline of defense activation, thus releasing them from reliance upon the microbiome. These results are particularly relevant to understanding the role that domestication, often involving polyploidization, has played in altering interactions between plants and their associated microbes in agricultural settings. Likewise, the protective effects of the SynCom in diploid plants have important implications on the role of Phyllosphere bacterial communities in managing plant disease, both naturally and as an applied supplement. Finally, researchers should be cautious about interpreting a lack of differences in microbiome composition or diversity as a lack of important differences between ploidy levels as evinced by our study.

## Methods

### *Arabidopsis* accessions

We used 14 total lines from 7 *Arabidopsis* diploids accessions from natural populations and their colchicine induced autotetraploids: Columbia (Col-0), Warschau (Wa-1), Wassilewskija (Ws-2), Gudow (Gd-1), HR (HR-5), Sorbo (Sorbo), St. Maria d. Feiria (Fei-0). We received Col-0 from Adrienne Roeder’s lab at Cornell, and Gd-1, HR-5, Sorbo, Fei-0 from Brian Husband’s lab at U. Guelph.

### Plant Growth Conditions

Seeds were surface sterilized by treatment with 70% ethanol for 2 min and then sodium hypochlorite solution (7% available chlorine) containing 0.2% Triton X-100 for 8 min. Samples were then washed seven times with sterile double distilled H2O (Bhardwaj et al., 2011). Seeds were then placed on MS media with .8% agar and cold stratify for 2 to 3 days at 4C in the dark (Bhardwaj et al., 2011). After germination, seedlings were transferred to a controlled environment with a long-day photoperiod (16-h photoperiod) at 22C and 55% relative humidity with cool white fluorescent light (Bhardwaj et al., 2011). After seven days the seedlings sprouts were transferred to sterile peat and the lighting was changed to short-day conditions (9-h photoperiod) (Innerebner et al., 2011).

### Inoculation with synthetic community (SynCom) and infection with pathogen DC3000

The synthetic community is composed of 25 taxa that span the diversity of microbial variation in tomato (Supplemental Table 1 Two weeks after germination, each plant was inoculated with either the synthetic community suspended in 10 mM MgCl_2_ buffer or just the 10 mM MgCl_2_ buffer as a control. The plants were inoculated by spraying the plant until saturation. Three weeks after germination (one week post synthetic community inoculation), the plants were spray-inoculated with either the pathogen (*Pseudomonas syringae* pv. tomato DC3000) or a 10 mM MgCl_2_ buffer. The pathogen inoculation was at a density of .0001 at OD600 (Innerebner et al., 2011).

### Sample Collection

Four sets of samples were collected, the first set one week post inoculation with the SynCom, but immediately prior to inoculation with DC3000 in order to determine the commensal community composition, the second immediately after inoculation with DC3000, the third 24 hours post inoculation, and the fourth 48 hours post inoculation. All of the plants were approximately at the same stage of development and no plants that showed signs of inflorescence emergence were used in the assay. To sample the aerial portion of the plants we cut just above the roots and transferred the total aboveground biomass into a tube with either 10 mM MgCl_2_ (for sequencing the SynCom), or into 100mM phosphate buffer (pH 7), for the pathogen inoculated samples. Samples for sequencing were sonicated for 15 minutes in a Branson M5800 sonicating water bath. The resulting leaf wash was then pelleted, the supernatant removed, and frozen at -20ºC until sequencing. Pathogen inoculated samples were bead homogenized using the FastPrep-24 Classic bead beating grinder and lysis system (MP Biomedicals, Inc., CA, USA) and frozen at -20ºC until ddPCR sequencing was performed.

### Amplification and Sequencing of Microbial 16S rDNA

Samples were frozen and kept at -20ºC and sent out to Microbiome Insights for 16S V4 sequencing and qPCR analysis within one month of freezing. Amplification and sequencing was performed according to Microbiome Insights standard protocol: Specimens were placed into a MoBio PowerMag Soil DNA Isolation Bead Plate. DNA was extracted following MoBio’s instructions on a KingFisher robot. Bacterial 16S rRNA genes were PCR-amplified with dual-barcoded primers targeting the V4 region (515F 5’-GTGCCAGCMGCCGCGGTAA-3’, and 806R 5’-GGACTACHVGGGTWTCTAAT-3’), as per the protocol of Kozich et al. (2013). Amplicons were sequenced with an Illumina MiSeq using the 300-bp paired-end kit (v.3). The potential for contamination was addressed by co-sequencing DNA amplified from specimens and from template-free controls (negative control) and extraction kit reagents processed the same way as the specimens. A positive control from ‘S00Z1-’ samples consisting of cloned SUP05 DNA, was also included. The only modification to this standard protocol was the addition of PNAs according to the method developed in Lundberg et al. 2012, in brief (mPNA, to knock out mitochondria and pPNA to knock out chloroplast) into the PCR step during library prep at a concentration of 5uM per PNA. The PCR reaction was then modified with the addition of a PNA annealing step at 78 °C for 10s.

### Microbial abundance through qPCR

From the standard methods of Microbiome Insights: Bacterial-specific (300 nM 27F, 5’-AGAGTTTGATCCTGGCTCAG-3’) forward primers coupled to (300 nM 519R, 5’-ATTACCGCGGCTGCTGG-3’) reverse primers were used to amplify Bacterial 16S rRNA. 25 μl reactions using iQ SYBR Green Supermix (Bio-Rad) were run on Applied Biosystems StepOne Plus instrument in triplicate. For standards, full-length bacterial 16S rRNA gene was cloned into a pCR4-TOPO vector, with Kanomycin-Ampicillin resistance. The total plasmid fragment size is expected to be 5556 bp. A bacterial standard was prepared via. 10-fold serial dilutions, and the copies of 16S was determined by the following: Copy# = (DNA wt. x 6.02E23)/(Fragment Size x 660 × 1E9). Linear regression was used to determine copy numbers of samples, based on CT of standards. Reaction specificity was assessed using a melt curve from 55 oC to 95 oC, held at 0.5 oC increment for 1s.

### Microbiome Data Analysis

Forward and reverse paired-end reads were filtered and trimmed to 230 and 160 base pairs (bps), respectively using the DADA2 pipeline with default parameters (Callahan et al., 2016). Following denoising and merging reads and removing chimeras (Table 2), we used DADA2 to infer amplicon sequence variants (ASVs) which are analogous to operational taxonomic units (OTUs) and taxonomy was assigned to these ASVs using the DADA2-trained SILVA database (Version 132, https://benjjneb.github.io/dada2/training.html). Using the negative samples from 16s sequencing we implemented the decontam package using default settings to identify and remove potential contamination from the samples (Davis et al. 2017). The assigned ASVs, read count data, and sample metadata were combined in a phyloseq object (McMurdie and Holmes, 2013) for downstream analyses. Differential microbial changes were calculated using DESeq2 (Love et al., 2014) and the phyloseq (Ssekagiri et al., 2018) package was implemented in R to calculate changes in alpha and beta diversity. For a permutational analysis of variance (PERMANOVA), data was rarified to 90% of the reads of the least abundant sample and the test was performed using the adonis function in the vegan package (v2.5-2, Oksanen et al. (2007)) in R with 999 permutations to test whether ploidy, or genotype had an effect on beta diversity measures.

### ddPCR pathogen assay

Absolute bacterial abundance was estimated by performing digital droplet PBR (ddPCR) on homogenized whole plant samples randomized within plate columns using the BIO-RAD QX 200 Droplet Reader (Bio-Rad Laboratories, Inc., Hercules, CA, USA) and custom primers to specifically target and amplify *Pseudomonas syringae* pv. tomato DC3000 (Supplemental Table 1). The PCR protocol is as follows: 95° for 5 min., 95° for 30 sec., 60° for 100 sec., return to step two 40 times., 4° for 5 min., 90° for 5 min., keep at 4° overnight. We used the default thresholds for identifying positive samples on the Biorad analysis software, and then used the weight of each sample to calculate a normalized per gram density of bacteria present on the above-ground plant. We compared the absolute abundance of polyploids and diploid accession pairs across each time point in order to assay how the pathogen interacted with ploidy and microbiome treatment.

### RNA Sample Collection and Sequencing

For each of three accessions (Columbia (Col-0), Wassilewskija (Ws-2), Sorbo (Sorbo)), we grew randomized blocks six plants of each ploidy level (diploids and induced autotetraploids) with three plants treated with the synthetic community and three treated with the control buffer, for a total of 36 plants. We collected single leaves from the largest developmental node of plants at Stage 1.10 (ten rosette leaves >1 mm in length (Boyes et al., 2001)) and directly froze them in liquid nitrogen before subsequent storage at -80C. Tissue was homogenized using a Mini-BeadBeater 8 (BioSpec Products, Bartlesville, OK, USA) following the manufacturer’s instructions. RNA was extracted using the Spectrum Plant Total RNA Kit (Merck / MilliporeSigma, MO, USA) according to the manufacturer’s recommendations. We pooled three samples per accession for three accessions and their synthetic autotetraploids. Samples were sent to Novogene USA Inc. (Sacramento, CA) for library prep (Poly(A) capture, ligation-based addition of adapters and indexes) and sequencing (Illumina NovaSeq 6000, paired-end reads of length 150 bps, 20M reads per sample).

### RNA-seq Data Processing and Analysis

Raw FASTQ files were trimmed and filtered to remove low-quality reads and technical sequences using Trimmomatic (Bolger et al., 2014) with the default settings. Filtered reads were aligned to the *Arabidopsis* reference sequence (TAIR10, Lamesch et al. (2012)) with HISAT2 (Pertea et al., 2016). HTSeq (Kim et al., 2015) was used to determine read counts per gene for the test for euploidy and DESeq2 was used to analyse differential gene expression (Love et al., 2014) for different experimental comparisons. For DESeq2 analysis gene ontology was assigned using UniProt (Ruch P et al., 2021). Links to the DESeq2 output data for each of these comparisons can be found in Supplemental Table 2. Enriched gene ontology (GO) terms were then identified using GOrilla (Eden et al., 2009). Further analysis was performed using iDEP: an integrated web application for differential expression and pathway analysis of RNA-Seq (Ge et al., 2018) data in order to assess patterns of differential gene expression and enrichment within Kyoto Encyclopedia of Genes and Genomes (KEGG) pathways (Kanehisa et al., 2017).

### Test for euploidy

We tested if the tetraploid samples were aneuploid or euploid by calculating fold change in relative expression (transcripts per million; TPM) per gene for every pairwise comparison of biological replicates following the methods outlined in Song et al. (2020). If there is aneuploidy, we would expect to see a large coordinated increase or decrease in TPM for genes on that chromosome, which would be reflected in a shift in fold change of expression relative to the other biological replicates (Supplemental Figure 1). We did not find any shift and therefore conclude that all tetraploid individuals were euploid.

## Acknowledgements

Work was supported by the National Science Foundation under Grant 1838299, as well as the Lawrence R. Heckard Endowment Fund of the Jepson Herbarium and an American Society of Plant Taxonomists W. Hardy Eshbaugh Graduate Student Research Grant Award to MJS.

## Supplemental Figures and Tables

**Supplemental Figure 1.**
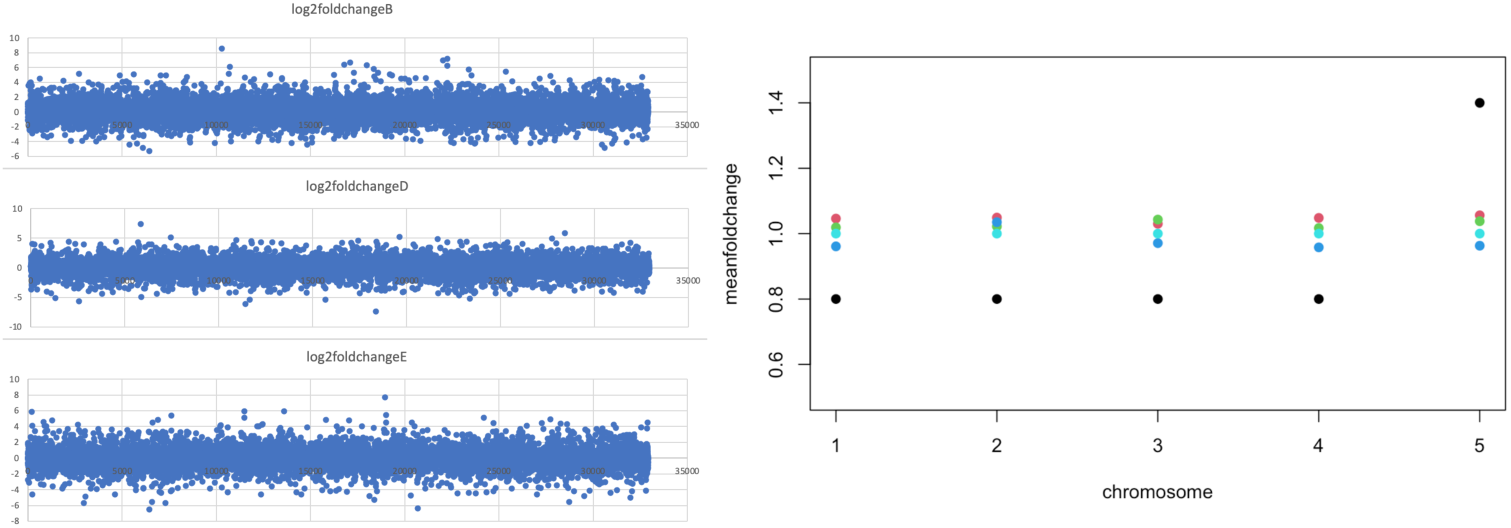
Assessment of polyploid lines for aneuploidy. Left: Transcripts per million (TPM) per gene among the biological replicates of an accession (Col-0 (top), Ws-2 (middle), or Sorbo (bottom)) and plotted along the length of all five chromosomes. If any one showed a stretch (or whole chromosome) of elevated or lowered TPM relative to any of the others it would suggest aneuploidy (chromosomal or segmental). Right: Blue, green, and red dots represent the mean fold change per gene per chromosome for Col-0, Sorbo, and Ws-2, respectively. Cyan dots represent the expected pattern for an euploid (all 1.0) and black dots represent the expected pattern for an aneuploid where there is a coordinated transcriptional increase due to a segmental or chromosome duplication.

**Supplemental Figure 2.**
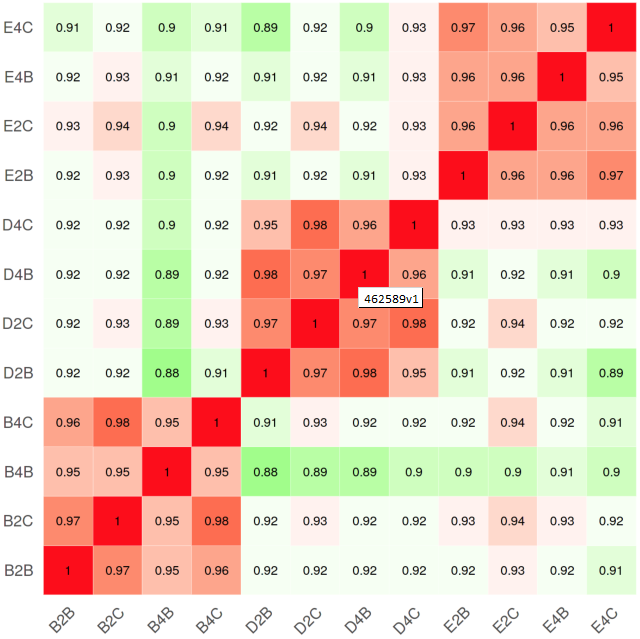
RNA seq heatmap of the sample-to-sample distances for Col-0 (B), Ws-2 (D), and Sorbo (E) accessions for treated (B) and untreated (C) plant samples.

**Supplemental Figure 3.**
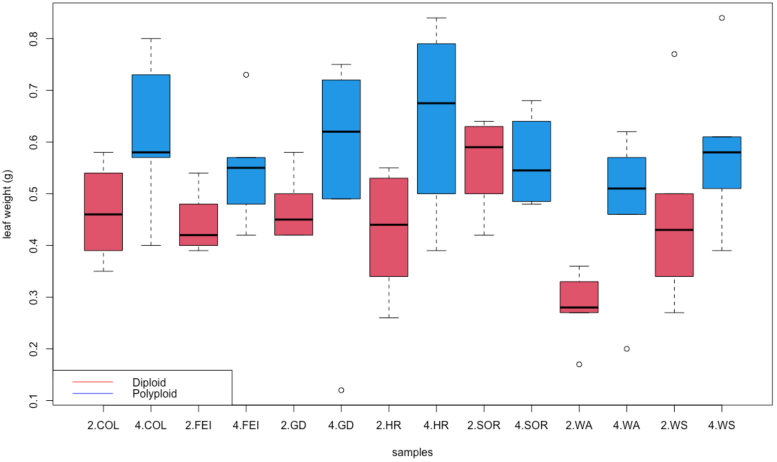
Leaf weights for diploid (red) and polyploid (blue) plants.

**Supplemental Figure 4.**
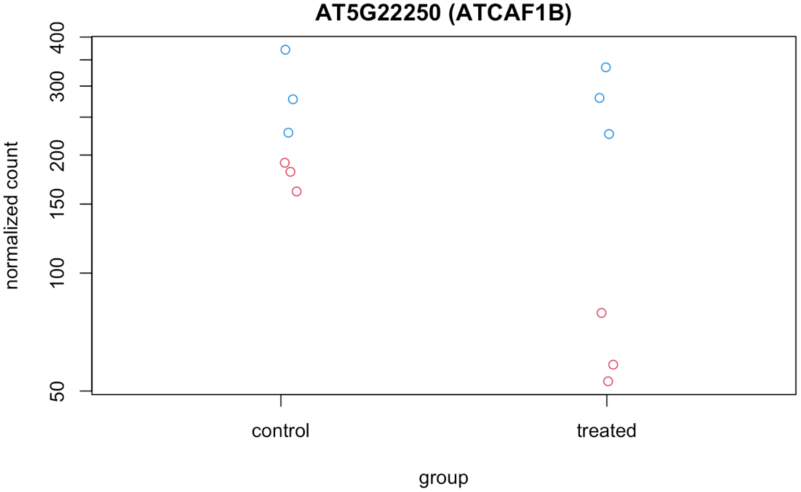
Normalized transcript counts for the AtCAF1B gene between diploids (red) and polyploids (blue) in microbiome treated and untreated samples.

**Supplemental Table 1.** Taxa List of Synthetic Community Members

| Genus | Species |
| --- | --- |
| *Brevibacterium* | *frigoritolerans* |
| *Bacillus* | *wiedmannii* |
| *Curtobacterium* | *herbarum* |
| *Curtobacterium* | *pusillum* |
| *Erwinia* | *tasmaniensis* |
| *Exiguobacterium* | *sibiricum* |
| *Frigoribacterium* | *endophyticum* |
| *Microbacterium* | *oleivorans* |
| *Pantoea* | *aurea* |
| *Pantoea* | *agglomerans* |
| *Pantoea* | *allii* |
| *Pseudomonas* | *asturiensis* |
| *Pseudomonas* | *rhizosphaerae* |
| *Pseudomonas* | *rhodesiae* |
| *Pseudomonas* | *moraviensis* |
| *Rathayibacter* | *festucae* |

**Supplemental Table 2.**
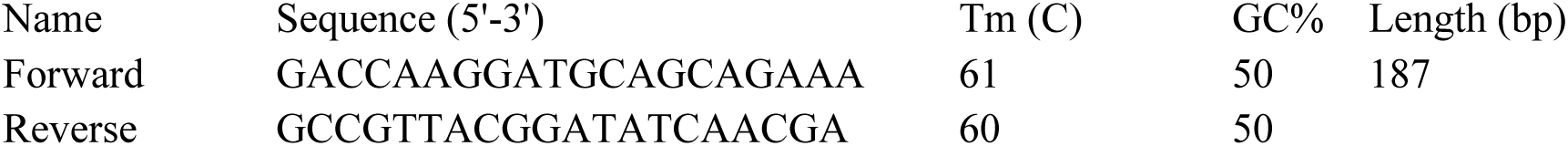
*Pseudomonas syringae* pv. tomato DC3000 specific primer used for ddPCR amplification

Supplemental File 1. Bacterial Abundance DESeq Results

Supplemental File 2. RNA DESeq Results

## References

1. Berg, M. and Koskella, B. (2018). Nutrient-and dose-dependent microbiome-mediated protection against a plant pathogen. Current Biology, 28(15):2487–2492.e3.

2. Bhardwaj, V., Meier, S., Petersen, L. N., Ingle, R. A., and Roden, L. C. (2011). Defence responses of Arabidopsis thaliana to infection by Pseudomonas syringae are regulated by the circadian clock. PloS One, 6(10).

3. Bodenhausen, N., Bortfeld-Miller, M., Ackermann, M., and Vorholt, J. A. (2014). A synthetic community approach reveals plant genotypes affecting the phyllosphere microbiota. PLoS Genet, 10(4):e1004283.

4. Bolger, A. M., Lohse, M., and Usadel, B. (2014). Trimmomatic: a flexible trimmer for illumina sequence data. Bioinformatics, 30(15):2114–2120.

5. Boyes, D. C., Zayed, A. M., Ascenzi, R., McCaskill, A. J., Hoffman, N. E.,Davis, K. R., and Gorlach, J. (2001). Growth stage–based phenotypic analysis of Arabidopsis: a model for high throughput functional genomics in plants. The Plant Cell, 13(7):1499–1510.

6. Bulgarelli, D., Schlaeppi, K., Spaepen, S., Van Themaat, E. V. L., and Schulze-Lefert, P. (2013). Structure and functions of the bacterial microbiota of plants. Annual Review of Plant Biology, 64:807–838.

7. Callahan, B. J., McMurdie, P. J., Rosen, M. J., Han, A. W., Johnson, A.J. A., and Holmes, S. P. (2016). Dada2: high-resolution sample inference from illumina amplicon data. Nature Methods, 13(7):581.

8. Castrillo, G., Teixeira, P. J. P. L., Paredes, S. H., Law, T. F., de Lorenzo, L., Feltcher, M. E., Finkel, O. M., Breakfield, N. W., Mieczkowski, P.,Jones, C. D., et al. (2017). Root microbiota drive direct integration of phosphate stress and immunity. Nature, 543(7646):513–518.

9. Chen, Z. J. (2010). Molecular mechanisms of polyploidy and hybrid vigor. Trends in Plant Science, 15(2):57–71.

10. Clay, N. K., Adio, A. M., Denoux, C., Jander, G., and Ausubel, F. M. (2009). Glucosinolate metabolites required for an Arabidopsis innate immune response. Science, 323(5910):95–101.

11. Coate, J. E., Luciano, A. K., Seralathan, V., Minchew, K. J., Owens, T. G., and Doyle, J. J. (2012). Anatomical, biochemical, and photosynthetic responses to recent allopolyploidy in Glycine dolichocarpa (Fabaceae). American Journal of Botany, 99(1):55–67.

12. Colaianni, N. R., Parys, K., Lee, H.-S., Conway, J. M., Kim, N. H., Edel-bacher, N., Mucyn, T. S., Madalinski, M., Law, T. F., Jones, C. D., andet al. (2021). A complex immune response to flagellin epitope variation in commensal communities. Cell Host & Microbe 29:635–649.e9.

13. Davis NM, Proctor D, Holmes SP, Relman DA, Callahan BJ (2017). Simple statistical identification and removal of contaminant sequences in marker-gene and metagenomics data. bioRxiv, 221499. doi: 10.1101/221499.

14. Ecker, J. R. and Davis, R. W. (1987). Plant defense genes are regulated by ethylene. Proceedings of the National Academy of Sciences of the United States of America, 84(15):5202–5206.

15. Eden, E., Navon, R., Steinfeld, I., Lipson, D., and Yakhini, Z. (2009). Gorilla: a tool for discovery and visualization of enriched go terms in ranked gene lists. BMC bioinformatics, 10(1):1–7.

16. Eulgem, T. and Somssich, I. E. (2007). Networks of wrky transcription factors in defense signaling. Current Opinion in Plant Biology, 10(4):366–371.

17. Felix, G., Duran, J. D., Volko, S., and Boller, T. (1999). Plants have a sensitive perception system for the most conserved domain of bacterial flagellin. The Plant Journal, 18(3):265–276.

18. Fukao, T. and Bailey-Serres, J. (2004). Plant responses to hypoxia–is survival a balancing act? Trends in Plant Science, 9(9):449–456.

19. Ge, S. X., Son, E. W., and Yao, R. (2018). iDep: an integrated web application for differential expression and pathway analysis of rna-seq data. BMC bioinformatics, 19(1):534.

20. Hijmans, R. J., Gavrilenko, T., Stephenson, S., Bamberg, J., Salas, A., and Spooner, D. M. (2007). Geographical and environmental range expansion through polyploidy in wild potatoes (Solanum section petota). Global Ecology and Biogeography, 16(4):485–495.

21. Huang, M.L., Deng, X.P., Zhao, Y.Z., Zhou, S.L., Inanaga, S., Yamada, S., and Tanaka, K. (2007). Water and nutrient use efficiency in diploid, tetraploid and hexaploid wheats. Journal of Integrative Plant Biology, 49(5):706–715.

22. Innerebner, G., Knief, C., and Vorholt, J. A. (2011). Protection of Arabidopsis thaliana against leaf-pathogenic Pseudomonas syringae by Sphingomonas strains in a controlled model system. Appl. Environ. Microbiol., 77(10):3202–3210.

23. Iqbal, N., Khan, N. A., Ferrante, A., Trivellini, A., Francini, A., and Khan, M. I. R. (2017). Ethylene role in plant growth, development and senescence: Interaction with other phytohormones. Frontiers in Plant Science, 8:475.

24. Kanehisa, M., Furumichi, M., Tanabe, M., Sato, Y., and Morishima, K. (2017). Kegg: new perspectives on genomes, pathways, diseases and drugs. Nucleic acids research, 45(D1):D353–D361.

25. Karasov, T. L., Chae, E., Herman, J. J., and Bergelson, J. (2017). Mechanisms to mitigate the trade-off between growth and defense. The Plant Cell, 29(4):666–680.

26. Kim, D., Langmead, B. and Salzberg, S.L., 2015. HISAT: a fast spliced aligner with low memory requirements. Nature methods, 12(4), pp.357–360.

27. King, K., Seppala, O., and Neiman, M. (2012). Is more better? polyploidy and parasite resistance. Biology Letters, 8(4):598–600.

28. King, K.C. and Bonsall, M.B., 2017. The evolutionary and coevolutionary consequences of defensive microbes for host-parasite interactions. BMC evolutionary biology, 17(1), pp.1-12.

29. Lamesch, P., Berardini, T. Z., Li, D., Swarbreck, D., Wilks, C., Sasidharan, R., Muller, R., Dreher, K., Alexander, D. L., Garcia-Hernandez, M., et al. (2012). The Arabidopsis information resource (TAIR): improved gene annotation and new tools. Nucleic Acids Research, 40(D1):D1202–D1210.

30. Lebeis, S. L., Paredes, S. H., Lundberg, D. S., Breakfield, N., Gehring, J., McDonald, M., Malfatti, S., Del Rio, T. G., Jones, C. D., Tringe, S. G., et al. (2015). Salicylic acid modulates colonization of the root microbiome by specific bacterial taxa. Science, 349(6250):860–864.

31. Leopold, D.R. and Busby, P.E., 2020. Host genotype and colonist arrival order jointly govern plant microbiome composition and function. Current Biology, 30(16), pp.3260-3266.

32. Levin, D. A. (1983). Polyploidy and novelty in flowering plants. The American Naturalist, 122(1):1–25.

33. Li, M., Xu, G., Xia, X., Wang, M., Yin, X., Zhang, B., Zhang, X., and Cui, Y. (2017). Deciphering the physiological and molecular mechanisms for copper tolerance in autotetraploid Arabidopsis. Plant Cell Reports, 36(10):1585–1597

34. Liang, W., Li, C., Liu, F., Jiang, H., Li, S., Sun, J., Wu, X., and Li, C. (2009). The Arabidopsis homologs of ccr4-associated factor 1 show mrna deadenylation activity and play a role in plant defence responses. Cell Research, 19(33):307–316.

35. Love, M. I., Huber, W., and Anders, S. (2014). Moderated estimation of fold change and dispersion for rna-seq data with deseq2. Genome Biology, 15(12):550.

36. Lundberg, D. S., Lebeis, S. L., Paredes, S. H., Yourstone, S., Gehring, J., Malfatti, S., Tremblay, J., Engelbrektson, A., Kunin, V., Del Rio, T. G., et al. (2012). Defining the core Arabidopsis thaliana root microbiome. Nature, 488(7409):86–90.

37. Mahalingam, R., Gomez-Buitrago, A., Eckardt, N., Shah, N., Guevara-Garcia, A., Day, P., Raina, R., and Fedoroff, N. V. (2003). Characterizing the stress/defense transcriptome of Arabidopsis. Genome biology, 4(3):R20.

38. Maier, B.A., Kiefer, P., Field, C.M., Hemmerle, L., Bortfeld-Miller, M., Emmenegger, B., Schäfer, M., Pfeilmeier, S., Sunagawa, S., Vogel, C.M. and Vorholt, J.A., 2021. A general non-self response as part of plant immunity. Nature Plants, 7(5), pp.696-705.

39. McLaren, M.R. and Callahan, B.J., 2020. Pathogen resistance may be the principal evolutionary advantage provided by the microbiome. Philosophical Transactions of the Royal Society B, 375(1808), p.20190592.

40. McMurdie, P. J. and Holmes, S. (2013). phyloseq: an R package for reproducible interactive analysis and graphics of microbiome census data. PloS One, 8(4).

41. Melotto, M., Underwood, W., Koczan, J., Nomura, K., and He, S. Y. (2006). Plant stomata function in innate immunity against bacterial invasion. Cell, 126(5):969–980.

42. Metcalf, C.J.E. and Koskella, B., 2019. Protective microbiomes can limit the evolution of host pathogen defense. Evolution Letters, 3(5), pp.534-543.

43. Molina-Henao, Y. F. and Hopkins, R. (2019). Autopolyploid lineage shows climatic niche expansion but not divergence in Arabidopsis arenosa. American Journal of Botany, 106(1):61–70.

44. Ng, D. W., Zhang, C., Miller, M., Shen, Z., Briggs, S., and Chen, Z. (2012). Proteomic divergence in Arabidopsis autopolyploids and allopolyploids and their progenitors. Heredity, 108(4):419–430.

45. Ni, Z., Kim, E.-D., Ha, M., Lackey, E., Liu, J., Zhang, Y., Sun, Q., and Chen, Z. J. (2009). Altered circadian rhythms regulate growth vigour in hybrids and allopolyploids. Nature, 457(7227):327–331.

46. Nuismer, S. L. and Thompson, J. N. (2001). Plant polyploidy and non-uniform effects on insect herbivores. Proceedings of the Royal Society of London. Series B: Biological Sciences, 268(1479):1937–1940.

47. Oksanen, J., Kindt, R., Legendre, P., O’Hara, B., Stevens, M. H. H., Oksanen, M. J., and Suggests, M. (2007). The vegan package. Community Ecology Package, 10:631–637.

48. Oswald, B. P. and Nuismer, S. L. (2007). Neopolyploidy and pathogen resistance. Proceedings of the Royal Society B: Biological Sciences, 274(1624):2393–2397.

49. Overmyer, K., Brosche, M., and Kangasjarvi, J. (2003). Reactive oxygen species and hormonal control of cell death. Trends in plant science, 8(7):335–342.

50. Pacey, E. K., Maherali, H., and Husband, B. C. (2020). The influence of experimentally induced polyploidy on the relationships between endopolyploidy and plant function in Arabidopsis thaliana. Ecology and Evolution, 10(1):198–216.

51. Pertea, M., Kim, D., Pertea, G.M., Leek, J.T. and Salzberg, S.L., 2016. Transcript-level expression analysis of RNA-seq experiments with HISAT, StringTie and Ballgown. Nature protocols, 11(9), pp.1650–1667.

52. Ponsford, J. C. B., Hubbard, C. J., Harrison, J. G., Maignien, L., Buerkle, C. A., and Weinig, C. (2020). Whole-genome duplication and host genotype affect rhizosphere microbial communities. bioRxiv, page 822726.

53. Ruch P, Teodoro D, UniProt Consortium. UniProt. 2021.

54. Saei, A., K. Hoeata, A. Krebs, P. Sutton, J. Herrick, M. Wood, and L. Gea. The status of Pseudomonas syringae pv. actinidiae (Psa) in the New Zealand kiwifruit breeding programme in relation to ploidy level. In IX International Symposium on Kiwifruit 1218, pp. 293–298. 2017.

55. Schoen, D., Burdon, J., and Brown, A. (1992). Resistance of Glycine tomentella to soybean leaf rust phakopsora pachyrhizi in relation to ploidy level and geographic distribution. Theoretical and Applied Genetics, 83(6-7):827–832.

56. Selosse, M.-A., Bessis, A., and Pozo, M. J. (2014). Microbial priming of plant and animal immunity: symbionts as developmental signals. Trends in Microbiology, 22(11):607–613.

57. Shi, H., Liu, W., Yao, Y., Wei, Y., and Chan, Z. (2017). Alcohol dehydrogenase 1 (adh1) confers both abiotic and biotic stress resistance in Arabidopsis. Plant Science, 262:24–31.

58. Song, M. J., Potter, B. I., Doyle, J. J., and Coate, J. E. (2020). Gene balance predicts transcriptional responses immediately following ploidy change in Arabidopsis thaliana. The Plant Cell, 32(5):1434–1448.

59. Ssekagiri, A., Sloan, W., and Ijaz, U. Z. (2018). microbiomeseq: an r package for microbial community analysis in an environmental context.

60. Tao, Y., Xie, Z., Chen, W., Glazebrook, J., Chang, H.-S., Han, B., Zhu, T., Zou, G., and Katagiri, F. (2003). Quantitative nature of Arabidopsis responses during compatible and incompatible interactions with the bacterial pathogen Pseudomonas syringae. The Plant Cell, 15(2):317–330.

61. Theodoridis, S., Randin, C., Broennimann, O., Patsiou, T., and Conti, E. (2013). Divergent and narrower climatic niches characterize polyploid species of european primroses in Primula sect. aleuritia. Journal of Biogeography, 40(7):1278–1289.

62. Ton, J., Flors, V., and Mauch-Mani, B. (2009). The multifaceted role of ABA in disease resistance. Trends in Plant Science, 14:310–317.

63. Valeri, M. C., Novi, G., Weits, D. A., Mensuali, A., Perata, P., and Loreti, E. (2020). Botrytis cinerea induces local hypoxia in Arabidopsis leaves. New Phytologist, 229: 173–185.

64. Van de Peer, Y., Mizrachi, E., and Marchal, K. (2017). The evolutionary significance of polyploidy. Nature Reviews Genetics, 18(7):411.

65. Vogel, C., Bodenhausen, N., Gruissem, W., and Vorholt, J. A. (2016). The Arabidopsis leaf transcriptome reveals distinct but also overlapping responses to colonization by phyllosphere commensals and pathogen infection with impact on plant health. New Phytologist, 212(1):192–207.

66. Walley, J. W., Kelley, D. R., Nestorova, G., Hirschberg, D. L., and Dehesh, K. (2010). Arabidopsis deadenylases atcaf1a and atcaf1b play overlapping and distinct roles in mediating environmental stress responses. Plant physiology, 152(2):866–875.

67. Wang, W., He, Y., Cao, Z. and Deng, Z., 2018. Induction of tetraploids in impatiens (Impatiens walleriana) and characterization of their changes in morphology and resistance to downy mildew. HortScience, 53(7), pp.925-931.

68. Wang, L., Cao, S., Wang, P., Lu, K., Song, Q., Zhao, F.-J., and Chen, Z. J. (2021). DNA hypomethylation in tetraploid rice potentiates stress-responsive gene expression for salt tolerance. Proceedings of the National Academy of Sciences, 118(13).

69. Wei, Z., Gu, Y., Friman, V.P., Kowalchuk, G.A., Xu, Y., Shen, Q. and Jousset, A., 2019. Initial soil microbiome composition and functioning predetermine future plant health. Science advances, 5(9), p.eaaw0759.

70. Wipf, H. M. and Coleman-Derr, D. (2021). Evaluating domestication and ploidy effects on the assembly of the wheat bacterial microbiome. Plos One, 16(3):e0248030.

71. Yasuda, M., Ishikawa, A., Jikumaru, Y., Seki, M., Umezawa, T., Asami, T., Maruyama-Nakashita, A., Kudo, T., Shinozaki, K., Yoshida, S., and et al. (2008). Antagonistic interaction between systemic acquired resistance and the abscisic acid–mediated abiotic stress response in Arabidopsis. 20:1678–1692.

72. Yoshida, T., Christmann, A., Yamaguchi-Shinozaki, K., Grill, E., and Fernie, A.R. (2019). Revisiting the Basal Role of ABA – Roles Outside of Stress. Trends in Plant Science 24, 625–635.

73. Zhang, X.-Y., Hu, C.-G., and Yao, J.-L. (2010). Tetraploidization of diploid Dioscorea results in activation of the antioxidant defense system and increased heat tolerance. Journal of plant physiology, 167(2):88–94.

74. Zhu, Q.-H., Stephen, S., Kazan, K., Jin, G., Fan, L., Taylor, J., Dennis, E. S., Helliwell, C. A., and Wang, M.-B. (2013). Characterization of the defense transcriptome responsive to fusarium oxysporum-infection in Arabidopsis using rna-seq. Gene, 512(2):259–266

